# Multivoxel Neural Reinforcement Changes Resting-State Functional Connectivity Within the Threat Regulation Network

**DOI:** 10.1101/2020.04.03.021956

**Authors:** Vincent Taschereau-Dumouchel, Toshinori Chiba, Ai Koizumi, Mitsuo Kawato, Hakwan Lau

## Abstract

Using neural reinforcement, participants can be trained to pair a reward with the activation of specific multivoxel patterns in their brains. In a double-blind placebo-controlled experiment, we previously showed that this intervention can decrease the physiological reactivity associated with naturally feared animals. However, the mechanisms behind the effect remain incompletely understood and its usefulness for treatment remains unclear. If the intervention fundamentally changed the brain responses, we might expect to observe relatively stable changes in the functional connectivity within the threat regulation network. To evaluate this possibility, we conducted functional magnetic resonance imaging (fMRI) sessions while subjects were at rest, before and after neural reinforcement, and quantified the changes in resting-state functional connectivity accordingly. Our results indicate that neural reinforcement increased the connectivity of prefrontal regulatory regions with the amygdala and the ventral temporal cortex (where the visual representations of phobic targets are). Surprisingly, we found no evidence of Hebbian-like learning during neural reinforcement, contrary to what one may expect based on previous neurofeedback studies. These results suggest that multivoxel neural reinforcement, also known as decoded neurofeedback (DecNef), may operate via unique mechanisms, distinct from those involved in conventional neurofeedback.

## Introduction

Exposure-based psychotherapies are amongst the most effective treatments for anxiety disorders (Craske et al., 2008). However, these interventions rely on the ability of patients to be exposed to the source of their fear which can create highly aversive reactions. As a result, many patients will terminate treatment prematurely (Loerinc et al., 2015) and some will even refuse to initiate treatment. This currently leaves a large number of patients without an adequate relief of their symptoms.

Advances in real-time fMRI (Shibata et al., 2011, 2019; Watanabe et al., 2017) allowed for the development of a new intervention that could be carried out without generating any aversive reaction (Koizumi et al., 2016; Taschereau-Dumouchel, Cortese, et al., 2018). This method called multivoxel neural reinforcement was designed according to the scientific principles of exposure-based psychotherapy (Craske et al., 2008). It involves pairing with a reward the unconscious activation of the multivoxel representation of a feared animal, such as a snake (see Figure 1a). We previously showed, in a double-blind placebo-controlled experiment, that this intervention can decrease physiological threat reactivity to feared animals -- as measured by amygdala and skin conductance reactivity.

**Figure 1.**
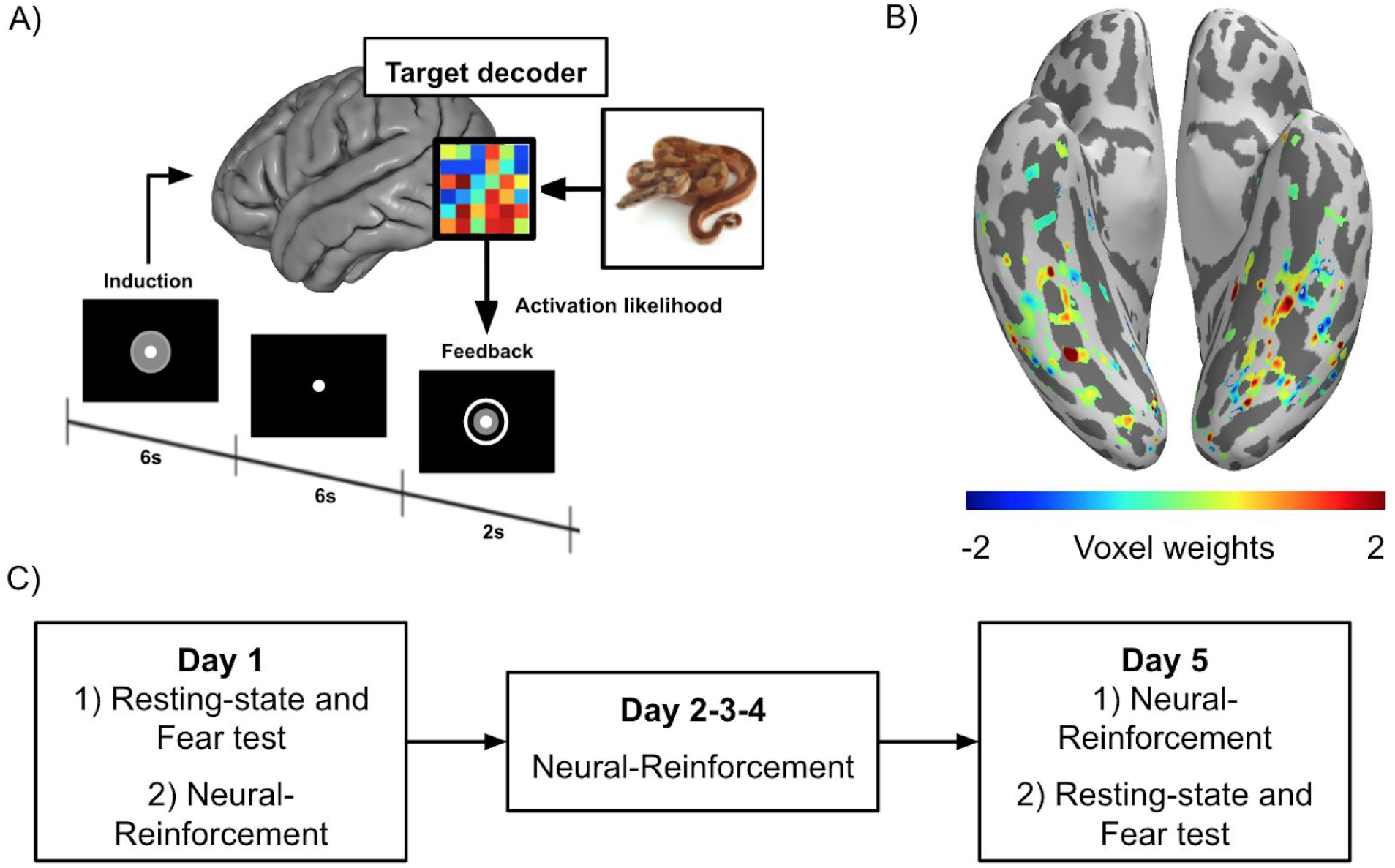
Summary of the neural reinforcement intervention (for more detailed information, see Taschereau-Dumouchel, Cortese, et al., 2018). A) Sequence of events in one trial of multivoxel neural reinforcement. During the induction period, the brain activity is processed online and decoded using the multivoxel representation of the target animal. This process provides us with an activation likelihood that is displayed visually to the participant. B) A representative multivoxel decoder of a target animal (voxel weights are standardised and slightly smoothed (FWHM = 1 mm) to facilitate the interpretation). These voxels were used as a seed region (which we call the ventral temporal cortex) in order to determine the changes in their connectivity following the intervention (The brain image was generated using pySurfer [https://pysurfer.github.io/]) C) Participants presenting self-reported fear of at least 2 animals included in our database took part in a neural reinforcement experiment. We used machine learning and a method called Hyperalignment (Haxby et al., 2011) in order to determine the multivoxel representation of the feared animals (i.e., decoders). The feared animal categories were then randomized to act either as the target or the control condition for the intervention. Participants completed five sessions of neural reinforcement conducted on different days. Before and after the intervention, participants completed resting-state sessions and were presented with images of the two animals they feared (i.e., fear test). During these sessions, participants were asked to report their subjective fear of the presented animals (The brain image was generated using Pycortex [Gao et al., 2015]).

However, many questions remain regarding the potential therapeutic applications of neural reinforcement. Notably, it is still unknown if multivoxel neural reinforcement can change the functional connectivity within the threat regulation network. Multiple previous studies reported such changes in functional connectivity following ROI-based (D. Scheinost et al., 2013) and functional connectivity neural reinforcement (Megumi et al., 2015; Yamashita et al., 2017). However, unlike most previous interventions that target specific regional activity, multivoxel approaches reinforce a distributed pattern of brain activity (see figure 1b). It is currently unknown if such sparse seed regions can also change their functional connectivity as a result of multivoxel neural reinforcement interventions.

Two key brain regions were primarily associated with our neural reinforcement procedure: the amygdala and the ventral temporal cortex (where the multivoxel representations of the feared animals were being reinforced during real-time fMRI). One possibility is that the amygdala and ventral temporal cortex might have changed their functional connectivity with some regulatory regions located in the prefrontal cortex. Two brain regions are primarily susceptible to be involved in this regulation. First, the ventromedial prefrontal cortex (vmPFC) has been shown to regulate amygdala reactivity during the extinction of threat memory (Phelps et al., 2004; Schiller et al., 2013) (however, see Fullana et al., 2018) and has been associated with the effect of a previous neural reinforcement intervention (Koizumi et al., 2016). Second, the ventrolateral prefrontal cortex (vlPFC) has also been shown to affect amygdala reactivity during threat exposure (Gold et al., 2015) as well as during emotion regulation (Braunstein et al., 2017; Morawetz et al., 2017; Silvers et al., 2016; Wager et al., 2008). Crucially, these regulatory regions have been associated with various anxiety disorders. For instance, social anxiety disorder has been associated with a decreased connectivity between the vmPFC and the ventral temporal cortex (Frick et al., 2013). Also, altered connectivity between the vlPFC and amygdala was associated with social anxiety disorders (Guyer et al., 2008) and generalized anxiety disorder (Monk et al., 2008). Therefore, investigating the changes in the regulatory activity of these regions might provide important insights to better understand the effects of neural reinforcement.

Another unanswered question regarding the potential therapeutic applications of neural reinforcement pertains to its capacity to change the subjective experience of fear. While physiological reactivity is undoubtedly an important mechanism of anxiety disorders (Etkin & Wager, 2007), there is also evidence suggesting that physiological reactivity might dissociate from the subjective fear reports (LeDoux & Brown, 2017; LeDoux & Pine, 2016; Taschereau-Dumouchel, Kawato, et al., 2019; Taschereau-Dumouchel, Liu, et al., 2018). Ultimately, successful clinical interventions should reduce subjective suffering. Therefore, future studies should continue to measure subjective ratings to assess the potential of the method for therapeutic developments.

To address these questions, we measured how the functional connectivity and subjective fear ratings changed as a function of the neural reinforcement intervention reported in Taschereau-Dumouchel and colleagues (2018). More precisely, in resting-state sessions, we determined if the amygdala and ventral temporal cortex changed their functional connectivity with the vlPFC and vmPFC following neural reinforcement. However, many other brain regions, such as the dorsomedial and dorsolateral prefrontal cortex, have also been associated with threat extinction and emotion regulation (Goodman, Harnett, and Knight 2018; Wager et al. 2008). Therefore, for the sake of transparency, we also conducted exploratory analyses investigating the changes in connectivity with all other cortical regions. Lastly, we investigated the changes in subjective fear ratings by presenting participants with pictures of the feared animals before and after neural reinforcement.

To anticipate, we found that neural reinforcement changed the resting-state functional connectivity between some key regions within the threat regulation network. However, these changes were not mirrored by any measurable decrease in the subjective fear reports. The theoretical and clinical implications of this dissociation will be discussed.

## Material and Methods

### Participants

17 participants (5 females; mean age = 21.92; SD = 1.54) took part in the neural reinforcement procedure previously described in Taschereau-Dumouchel et al., (2018). Participants were recruited if they reported “high” or “very high” levels of fear of at least 2 animals included in the original database. All participants provided written informed consent and the study was approved by the Institutional Review Board of Advanced Telecommunications Research Institute International, Japan.

### MRI parameters

Participants were scanned in two 3T MRI scanners (Prisma Siemens [20-channel head neck coil] and Verio Siemens [12-channel head coil]) at the ATR Brain Activation Imaging Center. During the experiments, we obtained 33 contiguous slices (TR = 2000 ms, TE =30 ms, voxel size = 3 × 3 × 3.5 mm ^3^, field-of-view = 192 × 192 mm, matrix size = 64 × 64, slice thickness = 3.5 mm, 0 mm slice gap, flip angle = 80 deg) oriented parallel to the AC-PC plane, which covered the entire brain. We also obtained T1-weighted MR images (MP-RAGE; 256 slices, TR = 2250 ms, TE = 3.06 ms, voxel size = 1 × 1 × 1 mm ^3^, field-of-view= 256 × 256 mm, matrix size = 256 × 256, slice thickness = 1 mm, 0 mm slice gap, TI = 900 ms, flip angle = 9 deg.).

### Procedure and processing of resting-state fMRI data

Resting-state fMRI data were acquired during two 8-minute sessions before and after neural reinforcement. Participants were asked to remain still while a fixation cross was projected on a translucent screen by an LCD projector (DLA-G150CL, Victor). The projector spanned 20 × 15 deg in visual angle (800 × 600 resolution) and had a refresh rate of 60 Hz. The fMRI images captured during the sessions were realigned to the first fMRI image, coregistered, motion-corrected (using 6 motion parameters), high-pass filtered (128-sec cutoff period), and normalized to the MNI space using SPM 12 (Statistical Parametric Mapping; www.fil.ion.ucl.ac.uk/spm) (Penny et al., 2011). To minimize saturation effects, the first 10 TRs were removed from further analysis. Linear detrending and scrubbing were conducted using inhouse functions in the Matlab R2014a environment (https://www.mathworks.com/products/matlab.html). Scrubbing was conducted such that the images presenting a frame displacement greater than 0.5 mm were removed from further analyses. The data were averaged by time point within each region of interest (see below for a definition of the regions) and standardised. The resulting time courses were correlated using partial correlations, where the global signal was introduced as a covariate. The obtained linear correlation coefficients were Fisher-transformed and compared between the two resting-state sessions (Post - Pre). Paired-sample t-tests were conducted to establish the significance of the change in connectivity. The effect sizes were calculated using Cohen’s d and corrected to account for the dependency between the means (S. B. Morris & DeShon, 2002). This was achieved using the compute_cohen_d function (https://www.mathworks.com/matlabcentral/fileexchange/62957-computecohen_d-x1-x2-varargin).

### The definition of the regions of interest

The masks of the amygdala and regulatory regions (i.e., vmPFC and vlPFC) were defined using the Brainnetome atlas, a connectivity based parcellation of the brain (Fan et al., 2016). In the absence of strong a priori hypotheses regarding laterality, the masks were created with the corresponding regions of both hemispheres. The amygdala mask regrouped all the amygdala subregions and was therefore generated by combining the medial and lateral regions defined by the Brainnetome atlas. The mask of the ventral temporal region was created using the voxels that presented significant weights during the neural reinforcement procedure (see (Taschereau-Dumouchel, Cortese, et al., 2018)). A representative ventral temporal mask is presented in Figure 1c. Since reinforcement sessions were conducted in the native space, the averaged activity of this mask was extracted in the native space.

With respect to the regulatory regions, both the vmPFC and vlPFC had many different definitions in the previous literature depending on the tasks (Buhle et al., 2014; Diekhof et al., 2011; Frank et al., 2014; Kalisch, 2009; Kohn et al., 2014; Messina et al., 2015) and populations studied (Dodhia et al., 2014; Etkin & Wager, 2007; Kircher et al., 2013; Klumpp et al., 2014; Sylvester et al., 2012; Young et al., 2017). Here the regions of interest were determined using previous meta-analyses (Fullana et al., 2016; Ipser et al., 2013) and the parcelation established by the Brainnetome atlas. As such, the vlPFC was defined as the inferior frontal sulcus (IFS) (Coordinates of the center of gravity in the MNI space: Left: [−47, 32, 14] and Right: [48, 35, 13]) and the vmPFC was defined as the medial area 14 (A14m) (Coordinates of the center of gravity in the MNI space: Left: [−7, 54, −7] and Right: [6, 47, −7]) and the medial area 10 (A10m) (Coordinates of the center of gravity in the MNI space: Left: [−8, 56, 15] and Right: [8, 58, 13]). Exploratory analyses were also conducted to investigate the changes in functional connectivity of the amygdala and ventral temporal cortex with all the other cortical regions defined by the Brainnetome atlas.

### Processing of the neural reinforcement fMRI data

We also aimed to determine how the activity within the ventral temporal region changed as a function of the neural reinforcement sessions. If the activity within the ventral temporal region globally increased or decreased as a function of neural reinforcement, this would suggest that Hebbian learning might have occurred between this region and other cortical regions. If so, this could explain the changes in resting-state functional connectivity. Therefore, we also conducted ROI analyses of the activity within the ventral temporal mask during the induction period of neural reinforcement using the structural masks of the ventral temporal region described above. The analyses were conducted using the same preprocessing procedure as the one previously used to compute neural reinforcement (see Taschereau-Dumouchel, Cortese, et al., 2018, 2019). One sample t-tests were conducted on the mean activity during each session of neural reinforcement.

### Subjective fear ratings

To assess changes in subjective fear ratings, we presented participants with images of animals from two feared categories (i.e., target and control categories from Taschereau-Dumouchel et al., 2018), These sessions were conducted before and after neural reinforcement (“Fear test” in Figure 1c). Each session included the presentation of 30 images divided in two short blocks: 10 images of the target condition, 10 images of the control condition, 5 images of a neutral animal (determined individually on a 7-point Likert scale), and 5 images of a neutral object. Each trial included the presentation of a fixation cross for 3–7 s (mean, 5 ± 2 s), presentation of the image for 6 s, and then a blank screen for 4–12 s (mean, 8 ± 3 s). Each block started with 20 s of rest, followed by the presentation of the image of a neutral object (e.g., a chair). The next two images were randomly set to be from the target or control category, and their order was counterbalanced between blocks. The remaining images were then presented randomly during the rest of the block. The subjective fear ratings were obtained by removing the mean fear rating of the neutral animal from the ratings of the feared animals (i.e., Rating of the feared animal minus the mean rating of the neutral animal). The subjective fear ratings were analyses using a 2×2 repeated-measure ANOVA conducted using the rm_anova2 function (https://www.mathworks.com/matlabcentral/fileexchange/6874-two-way-repeated-measures-anova)

## Results

### The connectivity of the amygdala

The results indicate that the connectivity between the amygdala and the vlPFC increased following the intervention (t(16) = 2.49; *P* = .024; Cohen’s d = 0.60). This change in connectivity was not observed between the amygdala and vmPFC (t(16) = 0.96; *P* = .35) (see Figure 2a and c).

**Figure 2.**
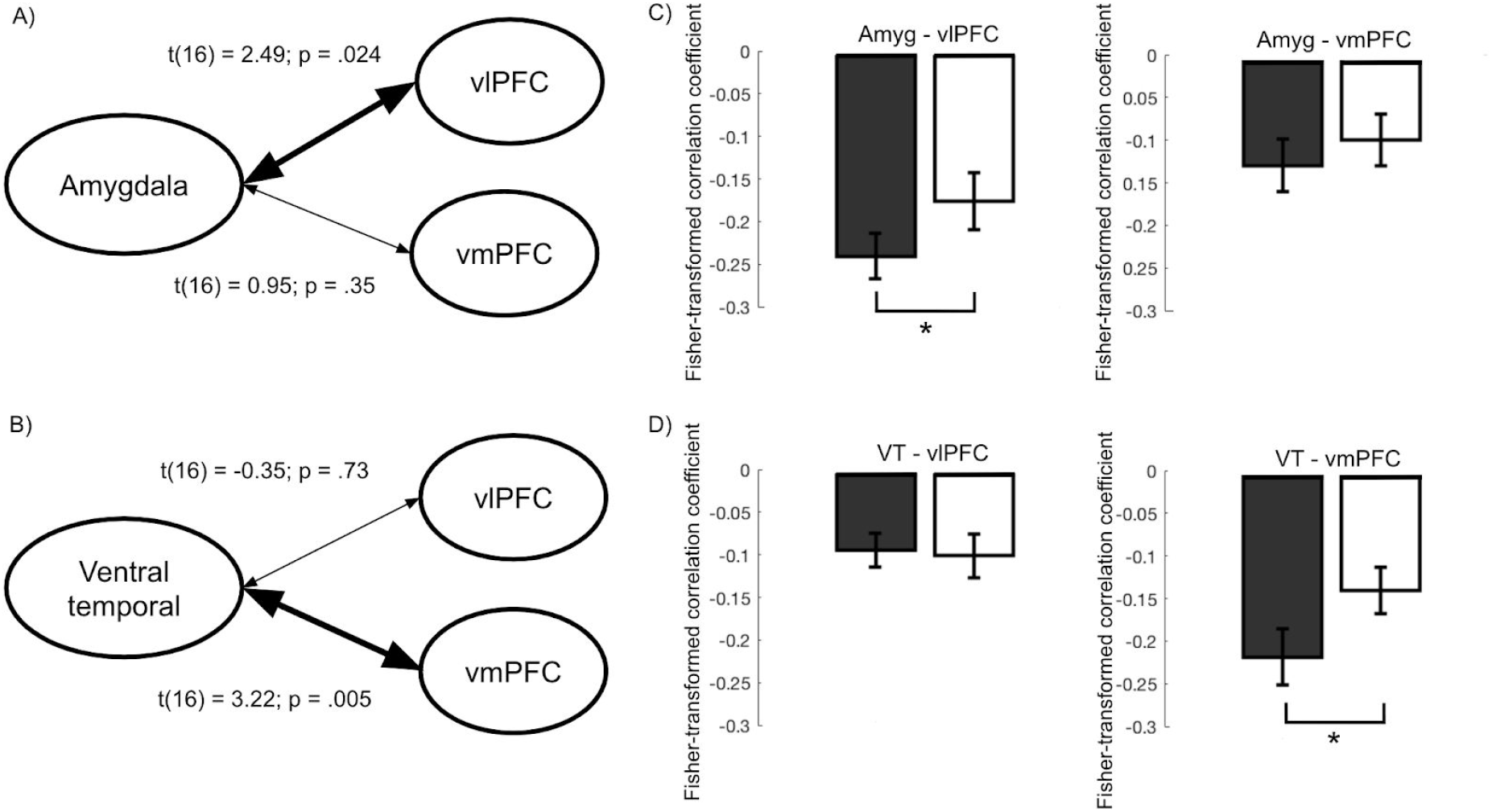
Changes in functional connectivity following neural reinforcement. Changes in connectivity (Post - Pre) between the regulatory regions (vlPFC and vmPFC) and (A) the amygdala and (B) the ventral temporal cortex. The Fisher-transformed correlation coefficients of (C) the amygdala and (D) the ventral temporal cortex before and after neural reinforcement. VT = Ventral temporal cortex; vlPFC = ventrolateral prefrontal cortex; vmPFC = ventromedial prefrontal cortex; Amyg = amygdala.

Exploratory analyses were also conducted regarding the changes in connectivity between the amygdala and the remaining cortical regions of the Brainnetome Atlas. The uncorrected results are presented in Figure 3a. These exploratory analyses revealed an increased in the correlation between the amygdala and the right hypergranular insula (t(16) = 2.64; *P* = .015; Cohen’s d = 0.66), the right cingulate cortex (subgenual area 32: t(16) = 2.30; *P* = .035; Cohen’s d = 0.56), the right fusiform gyrus (lateroventral area 37: t(16) = 2.69; *P* = .016; Cohen’s d = 0.63), and the left inferior parietal lobule (rostroventral area 40 (PFop); t(16) = 2.31; *P* = .035; Cohen’s d = .56). Furthermore, a decreased in the connectivity was also observed between the amygdala and the superior temporal gyrus (right caudal area 21: t(16) = −3.54; *P* = .0027; Cohen’s d = 0.86; Left rostral area 21: t(16) = −3.30; *P* = .0044; Cohen’s d = 0.80; Left anterior superior temporal sulcus: t(16) = −3.41; *P* = .0036; Cohen’s d = 0.83), the left fusiform gyrus (rostroventral area 20: t(16) = −2.80; *P* = .012; Cohen’s d = .68), the left inferior temporal gyrus (caudoventral area 20: t(16) = −2.22; *P* = .04; Cohen’s d = 0.54) and the left parahippocampal gyrus (temporal agranual insular cortex: t(16) = −2.41; *P* = −2.41; *P* = .028; Cohen’s d = .58).

**Figure 3.**
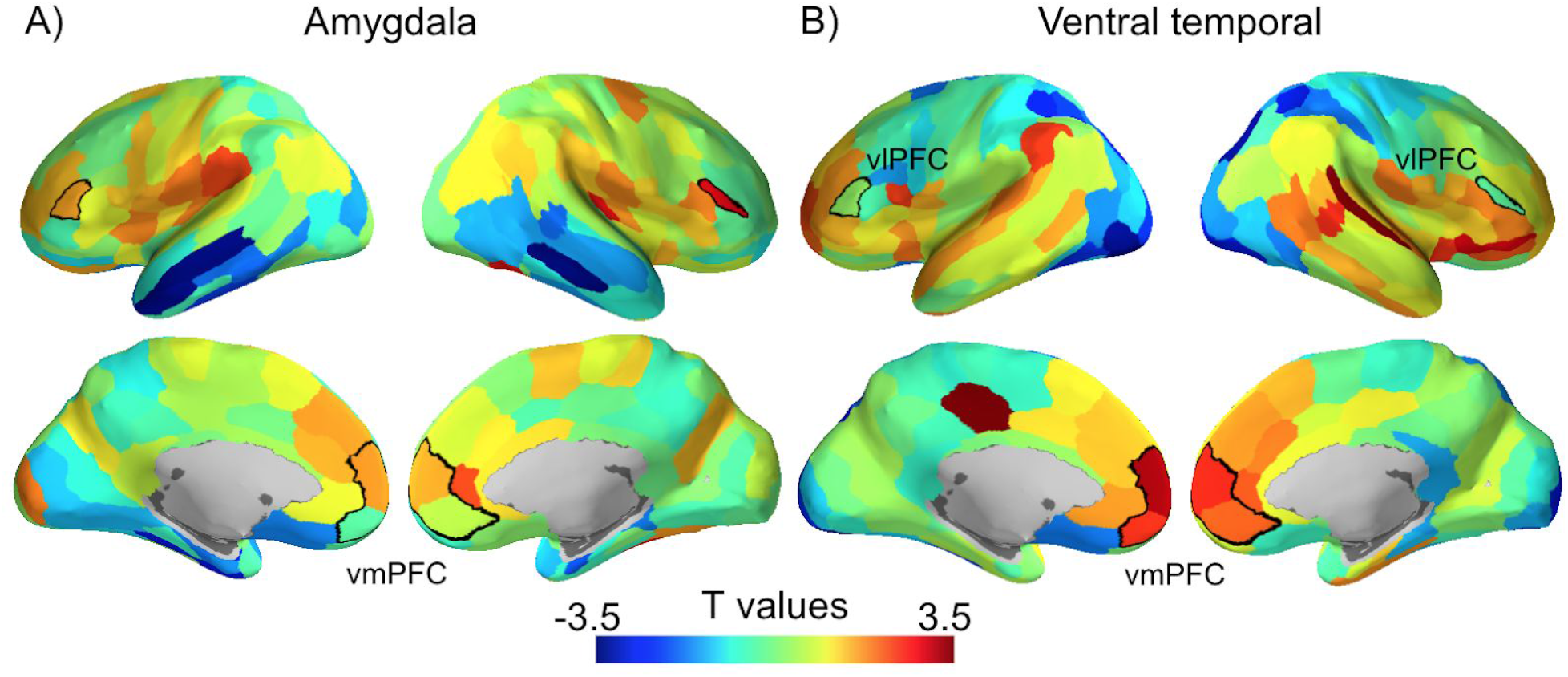
Exploratory analyses of the changes in connectivity of (A) the amygdala and (B) the ventral temporal cortex with the regions defined by the Brainnetome atlas. The *a priori* regulatory regions are bounded by a black line.

**Figure 4.**
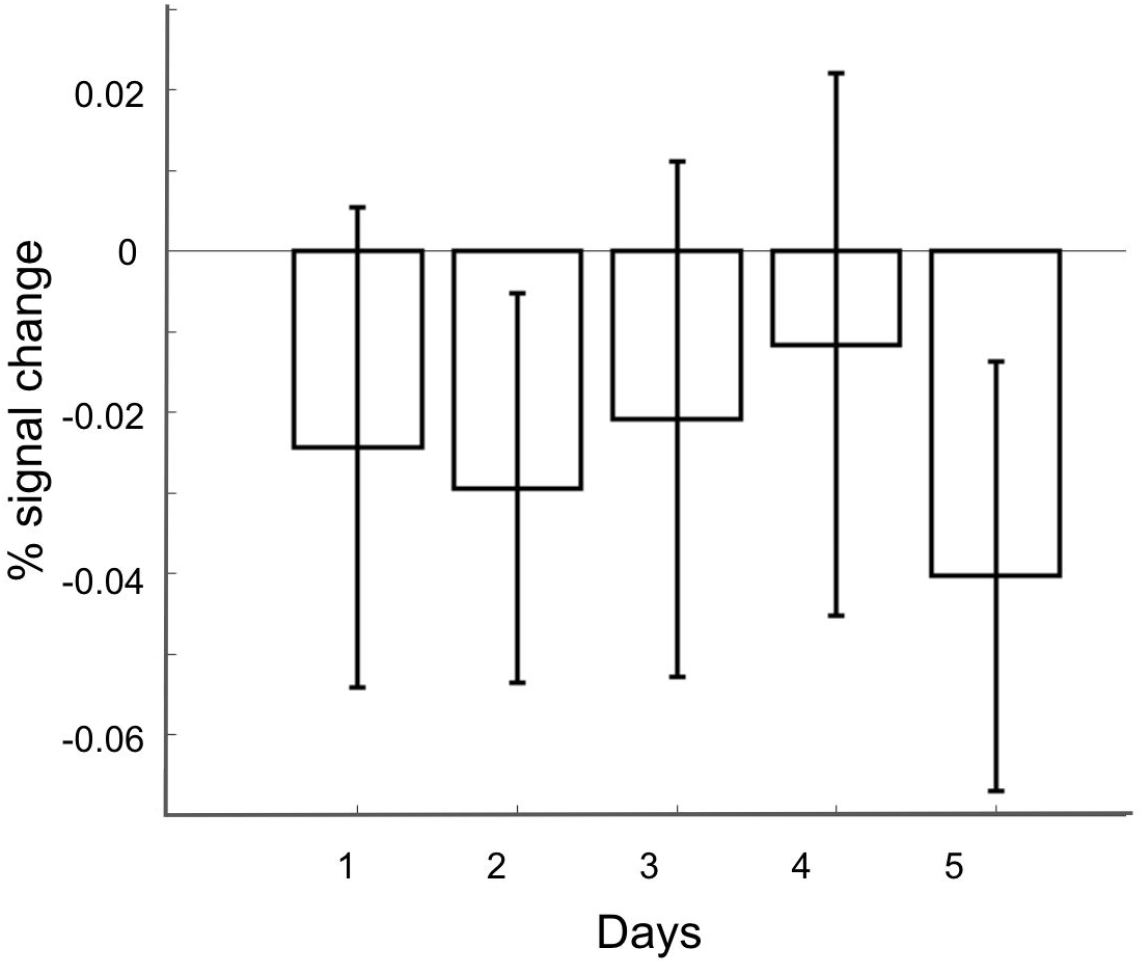
Percent signal change in the ventral temporal region during neural reinforcement. Signal within the ventral temporal region did not increase globally during neural reinforcement. These results suggest that Hebbian learning is unlikely to explain the changes in connectivity observed following the intervention.

### The connectivity of the ventral temporal cortex

The results indicate that the connectivity between the ventral temporal cortex and the vlPFC didn’t change following the intervention (t(16) = −0.35; *P* = .73) while the connectivity between the ventral temporal cortex and the vmPFC increased after neural reinforcement (t(16) = 3.22; *P* = .005; Cohen’s d = 0.78) (see Figure 2b and d).

Exploratory analyses were also conducted regarding the changes in connectivity between the ventral temporal cortex and the remaining cortical regions of the Brainnetome Atlas. The uncorrected results are presented in Figure 3b. These exploratory analyses revealed an increase in the connectivity between the ventral temporal cortex and the superior temporal gyrus (right caudal area 22: t(16) = 3.61; *P* = .0023; Cohen’s d = 0.88), the cingulate gyrus (left caudal area 23: t(16) = 3.47; *P* = .003; Cohen’s d = 0.84; right subgenual area 32: t(16) = 2.19; *P* = .04; Cohen’s d = 0.53), the orbital gyrus (right lateral area 12/47: t(16) = 2.90; *P* = .011; Cohen’s d = 0.70; right orbital area 12/47: t(16) = 2.18; *P =* .044), the posterior superior temporal sulcus (right rostroposterior superior temporal sulcus: t(16) = 2.54; *P* = .022; Cohen’s d = 0.62), insular gyrus (right ventral agranular insula: t(16) = 2.49; *P* = .024; 0.603; Cohen’s d = 0.60), inferior parietal lobule (left caudal areal 40 (PFop): t(16) = 2.431; *P* = .027; Cohen’s d = 0.59), inferior frontal gyrus (left ventral area 44: t(16) = 2.40; *P* = .029; Cohen’s d = 0.58), middle frontal gyrus (left area 46: t(16) = 2.257; *P* = .038; Cohen’s d = 0.55), Furthermore, these exploratory analyses revealed a decreased connectivity between the ventral temporal cortex and the superior parietal lobule (right intraparietal area 7 (hIP3): t(16) = −2.97; *P* = .009; Cohen’s d = 0.72; left intraparietal area 7 (hIP3): t(16) = −2.26; *P* = .038; Cohen’s d = 0.55; left lateral area 5: t(16) = −2.51; *P* = .023; Cohen’s d = 0.61), the lateral occipital cortex (right lateral superior occipital gyrus: t(16) = −3.41; *P* = .004; Cohen’s d = 0.83; left lateral superior occipital gyrus: t(16) = −2.82; *P* = .012; Cohen’s d = 0.68; left inferior occipital gyrus: t(16) = −3.30; *P* = .005; Cohen’s d = 0.80; right inferior occipital gyrus: t(16) = −2.78; *P* = .013; Cohen’s d = 0.67; right occipital polar cortex: t(16) = −2.63; *P* = .018; Cohen’s d = 0.64; left occipital polar cortex: t(16) = −2.33; *P* = .033; Cohen’s d = 0.57) and the fusiform gyrus (left lateroventral area 37: t(16) = −2.37; *P* = .031; Cohen’s d = 0.57).

### The activity of the ventral temporal region during neural reinforcement

The results indicate that the activity within the ventral temporal region did not increase or decrease during the induction period (All *P*s > .15).

### The subjective fear ratings

We conducted a repeated-measure ANOVA on the subjective fear ratings with one factor of Time (Pre and Post) and one factor of Conditions (Target and Control). The results indicate no effect of Time (F(1,16) = 0.14; *P* = .707; Partial Eta-Squared = .07) or Condition (F(1,16) = .091; *P* = .767; Partial Eta-Squared = .009) and no interaction effect (F(1,16) = .091; *P* = .767; Partial Eta-Squared = .006). More precisely, regarding the target of neural reinforcement, a direct comparison between the subjective fear rating before and after the intervention also indicated no effect of the intervention (t(16) = .449; *P* = .659; Cohen’s d = .10). The results are displayed in Figure 5.

**Figure 5.**
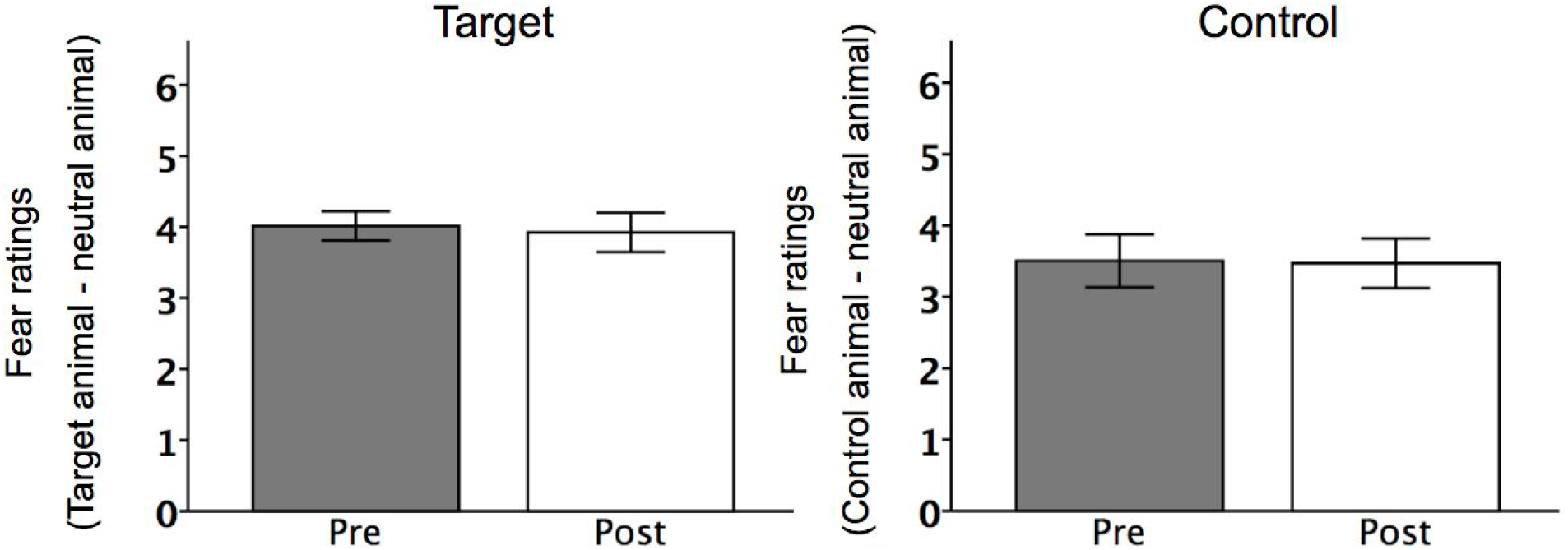
Neural reinforcement was not associated with a change in the subjective fear ratings. Participants presenting fear of at least two animal categories were recruited for the experiment. One of the feared category was randomly determined to be the target of the intervention while the other was determined to be the experimental control. The results indicate that, while participants presented a change in their physiological reactivity (see Taschereau-Dumouchel, Cortese, et al., 2018), their subjective fear ratings remained unchanged by the intervention.

## Discussion

Neural reinforcement has previously been shown to decrease physiological reactivity to feared animals (Taschereau-Dumouchel, Cortese, et al., 2018). In line with these findings, we here report that this intervention may also change the resting-state functional connectivity within the threat regulation network. More precisely, we showed that following the intervention, two key regions previously associated with neural reinforcement -- the amygdala and ventral temporal cortex -- increased their connectivity with the vlPFC and vmPFC, respectively. Importantly, these changes were not associated with any difference in the subjective fear ratings of participants. These results suggest the intriguing possibility that neural reinforcement might influence the threat regulation network unconsciously without changing the associated subjective experience -- at least not immediately.

This dissociation between threat reactivity and the subjective experience of fear has also been reported by multiple previous studies, primarily with respect to the activity of the amygdala. For instance, it has been reported that participants presenting congenital bilateral lesions of the amygdala could still experience fear under specific conditions (Anderson & Phelps, 2002; Feinstein et al., 2013). Furthermore, an important experimental demonstration recently showed that the direct electrical stimulation of the amygdala led only one out of nine patients to experience emotional reactions (Inman et al., 2018). These results are also in line with multiple experiments showing that threatening stimuli can generate amygdala reactivity even when participants are not consciously perceiving them (Morris et al., 1998, 1999; Ohman et al., 2007). As a result, some authors have called for caution in the direct association of implicit brain processes with subjective emotional states. This position is still controversial and is at the center of much recent debates and discussion (Fanselow & Pennington, 2017, 2018; LeDoux & Brown, 2017; LeDoux & Pine, 2016; Pine & LeDoux, 2017; Taschereau-Dumouchel, Liu, et al., 2018).

Based on these considerations, some may question the potential therapeutic applications of neural reinforcement. After all, a therapeutic intervention has to relieve the subjective suffering of patients in order to be clinically meaningful. However, few authors would actually suggest that the subjective experience of fear is totally unrelated to the underlying physiological reactivity. This view is notably supported by some recent computational models suggesting that the subjective experience most likely depends on a late-stage read-out of the sensory information (Maniscalco & Lau, 2016). If this is correct, then it might be possible to potentiate the current effects of Neural-Reinforcement. For instance, our intervention provided relatively little training to participants. Increasing the amount of neural reinforcement might strengthen the physiological changes and increase the probability to affect the subjective experience. Furthermore, combining neural reinforcement with conscious approaches such as cognitive restructuring (Craske & Barlow, 2006) might also help to decrease the subjective negative experience. This approach would be particularly interesting when exposure-based psychotherapies cannot be conducted with patients.

Another factor to consider is that the conscious effects of neural reinforcement might require some time to appear. For instance, it has recently been shown that the effects of neurofeedback can sometimes improve for weeks following the intervention (Rance et al., 2018). Furthermore, it is possible that following neural reinforcement multiple encounters with the phobic stimuli might be required for participants to observe the changes in their physiological reactivity and modify their interpretation of their experience. Therefore, it may be possible to potentiate the effects of neural reinforcement, even though no immediate changes of the subjective experience were observed yet.

Despite these considerations, we reported changes in resting-state functional connectivity that might be linked to many regulatory processes (Braunstein et al., 2017; Gyurak et al., 2011). For instance, the vlPFC has been associated both with implicit and explicit emotion regulation (Braunstein et al., 2017; Buhle et al., 2014). Also, the results of a meta-analysis revealed the role of the vlPFC in the success of cognitive-behavioral therapy for specific phobia (Ipser et al., 2013). Furthermore, while there is still some controversy regarding the precise role of vmPFC in emotion regulation (Fullana et al., 2018; Roy et al., 2012), multiple studies have reported its involvement in many regulatory processes, such as in the extinction of threat memories (Diekhof et al., 2011). Of a particular relevance, some results indicate that social anxiety disorders might be associated with a decreased connectivity between the vmPFC and the ventral temporal cortex (Frick et al., 2013). As such, our results could possibly reflect a normalization of some regulatory processes within the threat regulation network.

Admittedly, neural reinforcement is still a new intervention and few studies on its mechanisms of action are available (see (Oblak et al., 2017; Shibata et al., 2019). Our current results suggest the interesting possibility that the changes in the regulatory action of the vlPFC and vmPFC might have led to the decrease in amygdala and skin conductance reactivity. If this is the case, it might be interesting to consider how the baseline connectivity within the threat regulation network influences the success of such an intervention. If the functional connectivity within this network is indeed mechanistically related to the success of neural reinforcement, maybe this information can be exploited to determine which participants are most likely to benefit from neural reinforcement (see (Dustin Scheinost et al., 2014).

Hebbian-like learning is likely to explain changes in functional connectivity obtained following ROI-based (D. Scheinost et al., 2013) or functional connectivity neural reinforcement (Megumi et al., 2015; Yamashita et al., 2017). This is expected because these interventions aim to modulate altogether the activity of a specific brain region. However, multivoxel neural reinforcement targets a sparse group of voxels that can present both positive and negative relations to the activation likelihood. It is therefore surprising that this group of voxels presented a global change in their connectivity. These results open up a new question: what are the mechanisms of action that can explain such changes in connectivity? One may speculate that multivoxel neural reinforcement might have changed the expression of the target decoder during resting-state sessions. This might be a consequence of the increased activation likelihood observed during the neural reinforcement intervention (Shibata et al., 2011; Taschereau-Dumouchel, Cortese, et al., 2018). As such, studying how multivoxel neural reinforcement changes the expression of the target decoder during rest might ultimately help us to better understand the observed changes in functional connectivity.

One of the limitations of resting-state functional connectivity is that the observed changes cannot be directly linked to specific regulatory processes. Accordingly, here the regulatory effects of the vlPFC and vmPFC had to be inferred based on previous literature since participants were not asked to participate in a specific emotion regulation task. To better establish the association between changes in connectivity and the regulation of threat reactivity, it might be useful in the future to assess the regulatory processes more directly using experimental tasks (Braunstein et al., 2017; Gyurak et al., 2011).

Another concern is the small number of participants included in the study. Current evidence indicates that small samples are particularly problematic in fMRI as they can be severely underpowered (Button et al., 2013; David et al., 2013). Since neural reinforcement is a new approach, we first focused on providing a demonstration of its scientific principles. By increasing the number of participants, future studies might be able to provide more robust conclusions regarding the reported effects.

Finally, the duration of the effects of neural reinforcement still remains an open question. So far, we only investigated the changes in functional connectivity immediately after neural reinforcement. As discussed above, the effects of neurofeedback have been shown to increase over the weeks following the intervention (Rance et al., 2018). Future studies should determine the duration of the effects by including follow-up sessions in their experimental protocols. This appears to be important as it is one of the key concerns to address in order to determine the suitability of neural reinforcement for treatment.

## Conclusion

In sum, we showed that neural reinforcement might change the functional connectivity within the threat regulation network without changing the subjective experience of participants. While the mechanisms supporting this effect are still not well understood, this intriguing property of neural reinforcement might ultimately open new avenues for the treatment of anxiety disorders and help complement current therapeutic approaches.

## Acknowledgments

This research was conducted under the Japan Agency for Medical Research and Development (AMED) Grant Number JP18dm0307008. The study was also supported by the ImPACT Program of Council for Science, Technology and Innovation (Cabinet Office, Government of Japan). This work was partially supported by ‘Brain machine Interface Development’ under the Strategic Research Program for Brain Sciences supported by AMED. This study was also partly funded by the US National Institute of Neurological Disorders and Stroke of the National Institutes of Health (grant no. R01NS088628 to H.L.). V.T-D. is supported by a fellowship from the Fond de Recherche du Québec - Santé (FRQS) and a fellowship from the Canadian Institute of Health Research (CIHR). We thank K. Nakamura and M. Miuccio for their help in scheduling and conducting the experiment, N. Hiroe and H. Moriya for assistance with equipment and Y. Shimada and A. Nishikido for operating the fMRI scanner.

## Data availability

The data supporting the main findings of this manuscript are available from the ATR repository (https://bicr.atr.jp/decnefpro/). Matlab code can also be downloaded to recreate the statistical analyses and figures. These resources are available for non-commercial uses, and upon the reception of a signed agreement on terms and conditions for academic use.

